# The Kappa Opioid Receptor is required for some intermittent alcohol drinking induced changes in stress and threat responding in male C57BL/6J mice

**DOI:** 10.1101/2020.09.23.310284

**Authors:** Lara S. Hwa, Morgan Bowling, Rachel Calloway, Thomas L. Kash

## Abstract

The dynorphin/kappa opioid receptor (KOR) system in the brain regulates both stressful experiences and negative, aversive states during withdrawal from drugs of abuse. We explored the role of this system during acute withdrawal from long-term alcohol drinking. Male C57BL/6J mice were subjected to repeated forced swim tests, home cage exposure to a predator odor, and a visual threat after intermittent access to alcohol or water. Systemic injection of KOR antagonist norBNI reversed alcohol-related differences in immobility time during the second swim test and reduced burying behavior in response to predator odor, but did not affect behavioral response to visual threat.

**Highlights:**

- Intermittent alcohol drinking changed stress reactions in mice.
- KOR antagonist norBNI altered some, but not all, stress responses in alcohol drinkers

## 1. Introduction

The endogenous dynorphin/kappa opioid receptor (KOR) system is a regulator of stress responses and anxiety-like behavior (Bruchas, Land, and Chavkin 2010). Dynorphin/KOR signaling has also been heavily studied for treating negative symptomology observed in alcohol dependence (Walker et al. 2012; Kharkhanis et al. 2017). Rats rendered alcohol dependent with ethanol vapor show anxiety-like behavior during acute withdrawal, including decreased time spent in the open arm of the elevated plus maze, increased 22 kHz ultrasonic vocalizations, and increased marble burying, all of which can be blocked with the KOR antagonist norbinaltorphimine (norBNI, Valdez and Harshberger 2012, Berger et al. 2013, Rose et al. 2016). In this study, we sought to determine how alcohol drinking could impact multiple stress related behavioral responses and determine if the KOR was involved.

## 2. Materials and Methods

### 2.1 Animals

We used adult male C57BL/6J mice (Jackson Laboratories, Bar Harbor, ME) for behavioral pharmacology experiments with norBNI and Pdyn shRNA. Mice were singly housed in polycarbonate cages (GM500, Tecniplast, Italy) on a 12:12-h reversed dark-light cycle with lights off at 7:00am. Mice had *ad libitum* rodent chow (Prolab Isopro RMH 3000, LabDiet, St. Louis, MO) and H_2_O. The UNC School of Medicine Institutional Animal Care and Use Committee approved all protocols. Experiments were conducted in accordance with the NIH Guidelines for the Care and Use of Laboratory Animals.

### 2.2 Intermittent EtOH Drinking

Mice were given two-bottle choice intermittent access to a 20% (w/v) EtOH solution and water for at least 6 weeks (Hwa et al. 2011, Hwa et al. 2020). Drinking tubes positioned through customized cage lids consisted of conical centrifuge tubes (Catalog No. 05-502-10B; ThermoFisher) with rubber stopper (size 5.5, Ancare) and sippers (OT-101, 3.5 inch, Ancare). Bottles were weighed before and after 24 hour EtOH access on Mondays, Wednesdays, and Fridays. Fluid loss due to handling was averaged each week from a dummy cage and was subtracted from each mouse’s daily consumption. Mice were weighed to calculate daily EtOH intake in grams/kilogram (g/kg). Daily EtOH preference was defined as the ratio of EtOH fluid consumed divided by the total fluid (EtOH plus H_2_O) consumed.

### 2.3 Forced Swim Stress

After EtOH drinking, mice were exposed to two episodes of swim stress (SS) according to the protocol described by Anderson, Lopez, and Becker (2016). Mice were placed in 22-24°C water in a 20.32(w) x 26.67(h) cm acrylic cylinder for 10 min 4 hr before EtOH access. After each swim test, mice were dried and a heat lamp was kept over the home cage. There was one drinking day in between the double swim trial days (e.g. Monday and Friday swim stress preceding EtOH access on Monday, Wednesday, and Friday; **Fig 1A**). Latency to immobility and duration of immobility were recorded for both trials.

**Figure 1.**
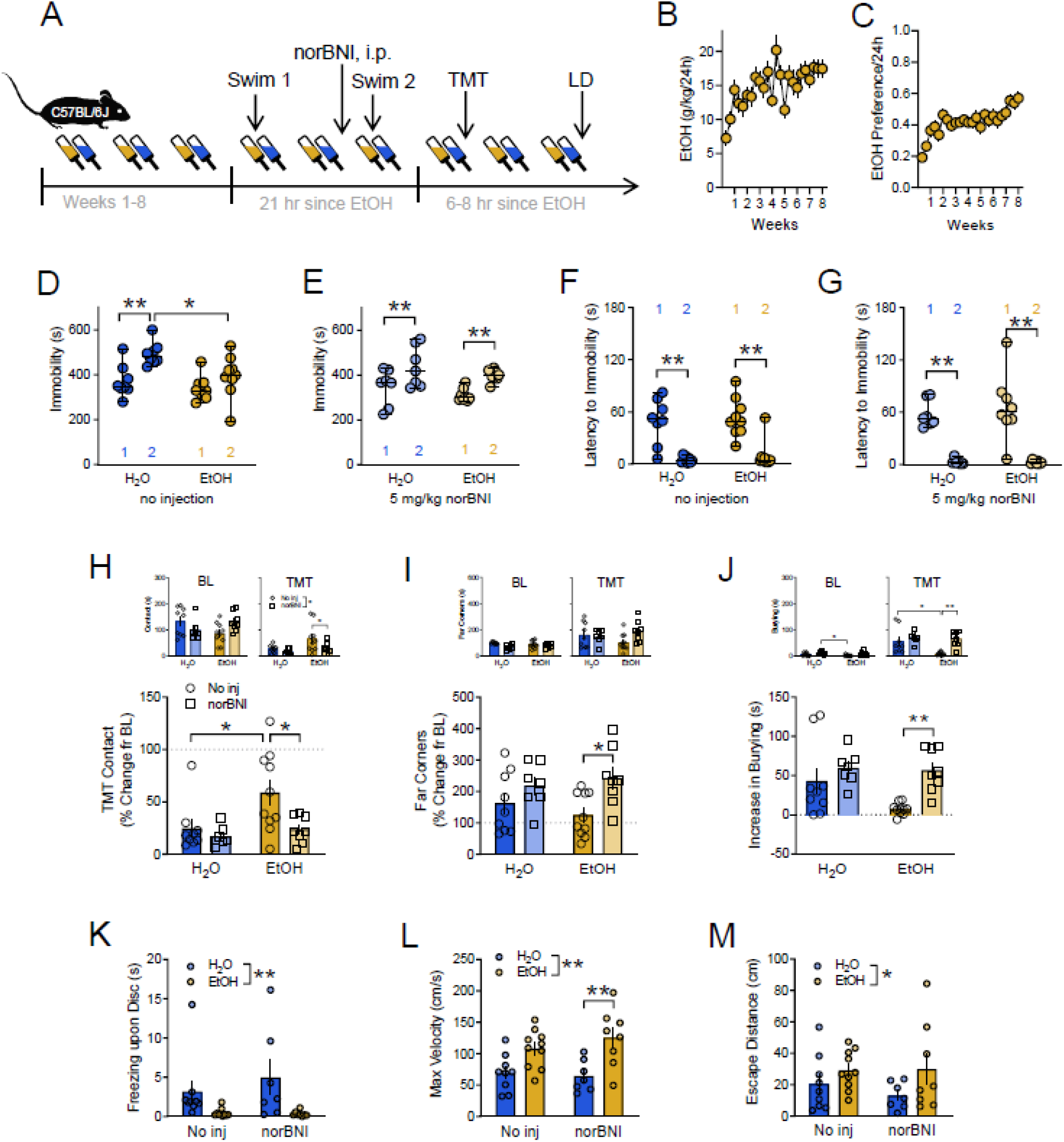
│Kappa opioid receptor regulation of alcohol-related responses to repeated swim test and predator odor. **A│** Experimental timeline of male C57BL/6J mice given two-bottle choice, intermittent EtOH for 8 weeks before stress testing in the repeated swim test, exposure to TMT predator odor, and an overhead looming disc. Long-lasting KOR antagonist norBNI was injected 16 hr before the second swim test. **B│** EtOH consumption (g/kg/24h) and **C│** EtOH preference ratio / 24 h across 8 weeks. **D│** Duration of immobility (s) across two forced swim trials in H_2_O-drinking (blue) and EtOH-drinking (gold) mice. Median ± range are displayed. **E│** Immobility (s) across repeated forced swim trials in norBNI-pretreated H_2_O (light blue) and EtOH (peach) mice. **F│** Latency to immobility (s) across swim trials and **G│** after norBNI treatment. In a home cage TMT exposure, percent change from baseline **H**│ contact time with the TMT (s), **I│** time spent in the far corners (s), and **J│** burying duration (s) in non-injected mice (circles) and norBNI-injected mice (squares). Mean ± SEM are displayed. Inset graphs are the raw durations (s) of behaviors during the 10 min baseline with the object (left panels) and during the 10 min TMT test (right panels). In reaction to the looming disc, **K**│ freezing behavior (s), **L**│ maximum velocity (cm/s) of the dart, and **M**│ escape distance (cm) to the hut were assessed in H_2_O (blue) and EtOH (gold) mice. *p<0.05. **p<0.01.

### 2.4 TMT Predator Odor

After repeated swim tests, mice were tested for their behavioral responses to the predator odor trimethylthiazoline (TMT) during acute, 6-8 hour, withdrawal from EtOH drinking [**Fig 1A**]. Testing occurred under 15-20 lux dim lighting conditions in a separate, ventilated room with a fume hood. After a 10 min habituation to a cotton tip applicator placed in the corner of the home cage (baseline, BL), 2.5 µl TMT (BioSRQ) was applied to the cotton tip followed by 10 min of behavioral testing. We have previously characterized that C57BL/6J male mice display a repertoire of stress-related and exploratory behaviors during the home cage TMT test, such as burying, freezing, grooming, rearing, stretch-attend, and walking (Hwa et al. 2020). Among these behaviors, burying represents a canonical stress or defensive-like behavioral state in rodents (Pinel and Treit 1978, De Boer and Koolhaas 2003). Durations of seconds 1) burying, 2) in contact with the TMT, and 3) in the far corners of the cage, were recorded and quantified with Ethovision XT13 (Noldus, The Netherlands). Percent change from the baseline was quantified for time spent in contact with the object and spent in the far corners, and raw increase in burying behavior in sec was measured from the pre-trial baseline.

### 2.5 Looming disc

After TMT exposure, mice were tested for reactions to an overhead looming disc during 6-8 hr withdrawal from intermittent EtOH [**Fig 1A**]. Based off of the protocol described by Yilmaz and Meister (2013), we used a custom open field chamber (48 x 48 x 30 cm) with three matte walls and one transparent wall for observation. One strip of red infrared LED lights provided ambient light in the test box, and an infrared camera (Basler GigE) recorded testing. A wide computer monitor (HP 22es, 48 cm) spanned across the test chamber to display the overhead looming disc animation. A triangular prism-shaped protective hut (12 x 20 cm) was positioned in the back corner during habituation and testing. Mice were placed into the test chamber for 10 min habituation to the arena, the protective hut, and the overhead monitor displaying a grey background (RGB 128, 128, 128). The looming disc animation was triggered when the mouse moved across the center of the arena after the 10 min habituation. The animation consisted of a 0.5 sec solid black circle rapidly increasing in size, repeated 15 times. In reaction to the looming disc, freezing behavior (s), maximum velocity (cm/s) of the dart, and the escape distance (cm) were recorded and quantified by Ethovision XT13.

### 2.6 NorBNI testing

Five mg/kg norBNI (Cat no. 0347, Tocris) was administered i.p., 1ml/100g, 16 hr before the second forced swim test to both EtOH and H_2_O mice. We have previously reported this time point for norBNI pretreatment to minimized handling stress prior to the stress reactivity testing and allowed for KOR antagonism (Hwa et al. 2020) instead of non-specific mu opioid antagonism that occurs initially post-injection (Endoh et al. 1992). Importantly, norBNI is known for its ultra-long duration of action (Munro et al. 2012). Since published reports have shown behavioral effects of norBNI for up to 86 days post-injection (Potter et al. 2011), it is likely that norBNI was on board during the second forced swim test, the TMT test, and the looming disc assay.

### 2.10. Statistical Analyses

Statistical tests were analyzed with GraphPad Prism 8 (La Jolla, CA, USA). Mixed model ANOVA was used to analyze immobility time and latency to immobility across Trials (Swim I and Swim II) between Drinking groups (H_2_O and EtOH). For TMT testing with norBNI or Pdyn-shRNA knockdown, two-way ANOVA was run to analyze contact with TMT, time spent in the far corners, and burying with Drug (no injection and norBNI) and Drinking as factors. Two-way ANOVA were similarly run for effects of norBNI on looming disc responses such as freezing, maximum velocity, and escape distance.

## 3. Results

### 3.1. Kappa opioid receptor regulation of alcohol-related responses to repeated swim test and predator odor

After a long-term history of voluntary EtOH drinking, mice were tested in three different stress and threat-related assays compared to single-housed, H_2_O-drinking controls [**Fig 1A**]. Adult male C57BL/6J mice escalated their daily alcohol consumption (g/kg) and alcohol preference on the intermittent, two-bottle choice schedule for 8 weeks [**Fig 1B-C**].

In the repeated forced swim test, H_2_O-drinking controls increased their immobility time from the first to the second swim, suggestive of learned stress coping responses, whereas EtOH drinkers did not [**Fig 1D**; Main effect of Trial F(1,15)=5.466, p=0.0337; Main effect of Drinking F(1,15)=22.61, p=0.003; Swim I H_2_O vs Swim II H_2_O t(15)=4.494, p=0.0009; Swim II H_2_O vs Swim II EtOH t(30)=2.933, p=0.0127]. In a separate group of mice, norBNI administration caused an increase in immobility time in the EtOH mice, implying a norBNI rescue of EtOH-impaired responding [**Fig 1E**; Main effect of Drinking F(1,13)=36.60, p<0.001; Swim I H_2_O vs Swim II H_2_O t(13)=4.170, p=0.0022; Swim I EtOH vs Swim II EtOH t(13)=4.398, p=0.0014]. Latency to immobility decreased from the first to the second swim [**Fig 1F**; Main effect of Trial F(1,15)=41.79, p<0.001; Swim I H_2_O vs Swim II H_2_O t(15)=4.489, p=0.0009; Swim I EtOH vs Swim II EtOH t(15)=4.663, p=0.0006]; however, EtOH history was not a factor. Latency to immobility was not affected by norBNI [**Fig 1G**; Main effect of Trial F(1,26)=59.29, p<0.001; Swim I H_2_O vs Swim II H_2_O t(26)=4.887, p<0.0001; Swim I EtOH vs Swim II EtOH t(26)=6.047, p<0.0001].

Next, mice were tested for their responses to TMT predator odor in the home cage. While TMT avoidance was maintained in H_2_O controls compared to non-odor baseline, norBNI decreased high levels of TMT contact in the EtOH mice [**Fig 1H**, Main effect of Drinking F(1,30)=5.950, p=0.0209; Main effect of Drug F(1,30)=5.438, p=0.0266; no injection H_2_O vs no injection EtOH t(30)=2.933, p=0.0377; no injection EtOH vs norBNI EtOH t(30)=2.919, p=0.0390]. Contact with the novel object during the no-odor baseline revealed an interaction between the factors, but post-hoc differences were not significant [**Fig 1H left inset**, Drinking x Drug Interaction F(1,30)=6.577, p=0.0156]. During the test, norBNI reduced time spent contacting the TMT in EtOH mice [**Fig 1H right inset**, Main effect of Drinking F(1,30)=4.255, p=0.0479; Main effect of Drug F(1,30)=5.530, p=0.0255; no injection EtOH vs norBNI EtOH t(30)=1.697, p=0.049]. NorBNI additionally increased percent change from baseline time spent in the far corners in EtOH mice [**Fig 1I**, Main effect of Drinking F(1,30)=8.880, p=0.0057; no injection EtOH vs norBNI EtOH t(30)=2.954, p=0.0357]. There were no EtOH or norBNI effects on raw time spent in the far corners in either the baseline period or the TMT test [**Fig 1I left and right inset**]. TMT-induced burying as a function of baseline increased with norBNI treatment in EtOH mice [**Fig 1J**, Main effect of Drinking F(1,30)=10.38, p=0.0031; no injection EtOH vs norBNI EtOH t(30)=3.491, p=0.0090]. Minimal burying activity during the baseline was different for norBNI-injected H_2_O mice and non-injected EtOH mice [**Fig 1J left inset**, Main effect of Drinking F(1,30)=8.811, p=0.0058; norBNI H_2_O vs no injection EtOH t(30)=3.192, p=0.0197]. Burying behavior in response to TMT was reduced in EtOH mice, which was restored in EtOH mice by norBNI pretreatment [**Fig 1J right inset**, Main effect of Drinking F(1,30)=12.06, p=0.0016; Main effect of Drug F(1,30)=5.950, p=0.0208; no injection H_2_O vs no injection EtOH t(30)=3.281, p=0.0156; no injection EtOH vs norBNI EtOH t(30)=3.932, p=0.0028].

Third, mice were assessed for reactions to the overhead looming disc. EtOH mice spent less time than controls freezing upon looming disc presentation, but norBNI did not affect these reactions in either group [**Fig 1K**, Main effect of Drinking F(1,30)=9.659, p=0.0041]. Further, EtOH mice had a heightened maximum velocity (cm/s) of the escape dart to the protective hut, but norBNI did not alter this response [**Fig 1L**, Main effect of Drinking F(1,30)=18.53, p=0.0002; norBNI H_2_O vs norBNI EtOH t(30)=3.538, p=0.0080]. The escape distance (cm) was overall farther for EtOH mice compared to H_2_O mice, but again, norBNI treatment was not a factor [**Fig 1M**, Main effect of Drinking F(1,30)=4.215, p=0.0489].

## 4. Discussion

Here, we found that KOR antagonist norBNI altered EtOH-induced disruptions in a repeated forced swim test and exposure to predator odor, but did not change reactions to a looming disc. Alcohol driven reductions in response to stress have been well-studied in humans (Sher 1987), so we aimed to determine how alcohol consumption could alter stress- and threat-related phenotypes in male C57BL/6J mice. In the forced swim test, EtOH and H_2_O mice showed no differences in immobility duration or latency to immobility in the forced swim test after eight weeks intermittent EtOH drinking. Others have also reported no differences in first swim test immobility between controls and EtOH mice after chronic intermittent EtOH vapor or ip EtOH injections at various withdrawal time points (Ribeiro-Carvalho et al. 2011, Bray et al. 2017, Maldonado-Devincci et al. 2016). There has been a resurgence in behavioral interpretation of the rodent forced swim test; besides anti-depressive-like behavior, immobility can also be interpreted as stress coping behavior, especially during repeated trials (Commons et al. 2017, Molendijk and De Kloet 2019). In agreement with this literature, we interpret the subsequent increase in immobility time during the second swim trial as a learned adaptive state, to conserve energy resources by staying immobile, which was evident in H_2_O controls but not in EtOH drinkers. This bears some superficial similarity to the immobility observed in H_2_O controls in response to the overhead looming disc. In contrast, EtOH mice exhibited much less immobility in favor of immediate darting behavior, suggestive of hyperresponsivity and differential assessment of the overhead threat. These reactions were accompanied by a longer escape path, which we additionally consider maladaptive in reaction to descending threat.

We also tested mice in their home cage for avoidance and burying responses to the predator odor TMT. EtOH mice generally showed increased contact with the TMT and reduced burying, which we have shown persists into protracted abstinence (Hwa et al. 2020). Defensive burying is a prominent stress coping strategy when rodents are confronted with noxious stimuli (De Boer and Koolhaas 2003), suggesting altered coping strategy. Taken together, these results suggest a history of EtOH drinking may alter stress and threat responses. Many others have observed EtOH withdrawal impairments in anxiety-like behavior in classical tests such as: deficits in social interactions (File et al. 1989, Lowery-Gionta et al. 2015, Marcinkiewcz et al. 2015), elevated plus maze (Lal et al. 1991; Wilson et al. 1998), elevated zero maze and light-dark test (Kliethermes et al. 2004) and novelty-induced hypophagia (Pang et al. 2013, Sidhu et al. 2018).

In addition, it was found that intermittent EtOH consumption led to increased sensitivity to social defeat stress (Nenning et al. 2020). These ‘affective disturbances’ during withdrawal (Holleran et al. 2016) may be also caused by slowed learning or low cognitive flexibility after intermittent EtOH exposure (Sey et al. 2019). Future studies should investigate stress-related cognitive impairment or behavioral flexibility in more formal tasks of decision making.

Comparing our TMT-evoked stress reactions to others, some have reported increased digging activity in 10-day withdrawal from chronic intermittent EtOH vapor (Sidhu et al. 2018), but this discrepancy may be explained by differential performance in the familiar home cage in response to TMT versus spontaneous digging activity in a novel cage. The Bains group, however, found decreased home cage digging after acute footshock stress (Füzesi et al. 2016), which is similar to the current results in EtOH-exposed mice after TMT. Our findings in the looming disc assay compliment the increased defensive responses to the bottle brush defined as increased escaping, which may be related to irritability or aggression found during withdrawal (Sidhu et al. 2018, Hwa et al. 2015.)

The present experiments have limitations. In the behavioral pharmacology studies, we acknowledge there is no vehicle injection group. As we were exploring subtle differences in stress behavior, we employed a non-injected, baseline behavioral strategy instead. Concerns for norBNI not being on board during stress testing are mitigated by the fact that systemic norBNI lasts for months post-injection (Potter et al. 2011).Another consideration worth discussing is the repeated stress tests. It is plausible that the repeated forced swim tests affected the TMT behavioral tests

## Abbreviations

(AUD): Alcohol use disorder
(EtOH): alcohol
(H_2_O): water
(Pdyn): preprodynorphin
(TMT): trimethylthiazoline
(SS): forced swim stress
(NS): non-stress
(KOR): kappa opioid receptor
(BL): baseline
(norBNI): norbinaltorphimine
(BNST): bed nucleus of the stria terminalis
(PBS): phosphate buffer solution
(TSA): tyramine signal amplification.

## Acknowledgements

This work was supported by the National Institutes of Health [grant numbers K99AA027576 (LSH) and U01AA020911 (TLK)

## Notes

### Competing Interest Statement

The authors have declared no competing interest.

### Summary of Updates

Several figures removed that had data with mistakes regarding animal sex (Figure 2 and 3). Figure 4 was removed because more recent validation of the pDYN shRNA construct in the BNST indicated it had only a small reduction on pDYN mRNA.

